# Using Published Pathway Figures in Enrichment Analysis and Machine Learning

**DOI:** 10.1101/2023.07.06.548037

**Authors:** Min-Gyoung Shin, Alexander R. Pico

## Abstract

Pathway Figure OCR (PFOCR) is a novel kind of pathway database approaching the breadth and depth of Gene Ontology while providing rich, mechanistic diagrams and direct literature support. PFOCR content is extracted from published pathway figures currently emerging at a rate of 1000 new pathways each month. Here, we compare the pathway information contained in PFOCR against popular pathway databases with respect to overall and disease-specific coverage. In addition to common pathways analysis use cases, we present two advanced case studies demonstrating unique advantages of PFOCR in terms of cancer subtype and grade prediction analyses.

## Background

For the past few decades pathway databases have relied on the manual extraction of pathway knowledge from the literature by teams of biocurators ranging from small, centralized groups to communities of hundreds of contributors [1–6]. Individuals comb the literature and contribute their domain knowledge in order to model the vast diversity of biochemical reactions and cellular processes comprising biological systems. Pathway databases are commonly used in enrichment analysis (a.k.a., pathway analysis), where they are reduced to collections of gene sets and tested for statistical overrepresentation or enrichment with respect to a researcher-provided gene set [7,8]. The connections and mechanistic details present in pathway models are still relevant for interpretation and visualization of enrichment results and are a key advantage of pathway databases over typical gene set collections such as GO. However, pathway databases also suffer disadvantages relative to simpler forms of gene set annotation, namely limited breadth, depth and curation throughput.

While GO Biological Process terms annotate 62% of human genes, pathway databases only cover up to 44% [9]. This limited breadth means that a significant percentage of a researcher’s genes of interest (e.g., from a differential expression dataset) would be essentially excluded from an enrichment analysis as they would yet be included in *any* pathway model. In terms of depth, pathway databases have traditionally focused on canonical pathways, for example having a single, generic representation of “apoptosis” or “hippo signaling”, despite the actual diversity of these biological processes across cell types, developmental stages, disease states and conditions. This oversimplification is understandable given the low throughput of pathway database content. Constructing a pathway model takes significant time and effort, including information gathering, synthesis, encoding and review [10,11]. Keeping up with the continuous flood of published findings spanning all aspects of cellular biology with manual curation is clearly a Sisyphean task. The WikiPathways project has had some success in addressing these challenges, sharing the burden of biocuration with any interested member of the research community [5,12]. Steadily acquiring ∼90 new pathways a year and ∼250 edits per month by a total of over 700 contributors, the WikiPathways project still pales in comparison to the volume of unique pathway diagrams routinely published in the literature as static images.

The Pathway Figure OCR (PFOCR) project takes a more direct approach to capturing pathway information in a pathway database [13]. Collecting 1,000 published pathway figures per month in recent years and a total of 79,949 pathway figures since 1995 from the indexes of PubMed Central, the project has extracted 1.5 million human genes (14,253 unique; 60% of all human genes), 218 thousand chemicals (11,100 unique), and 29 thousand disease names (1,204 unique) via a pipeline involving machine learning, optical character recognition (OCR), and named entity recognition (NER). The PFOCR database contains more unique genes than any other pathway database and is comparable in breadth to Gene Ontology. With remarkably little redundancy, PFOCR contains many dozens of unique instances of processes like apoptosis and hippo signaling that typically have only canonical representations in pathway databases. In terms of throughput, the algorithmic steps of the PFOCR project could feasibly generate an up-to-date database in lockstep with the indexing efforts of PubMed Central.

In this paper, we will provide a detailed characterization of the PFOCR collection of pathways with a particular focus on pathways and genes with disease associations. We will also demonstrate the utility of PFOCR in comparison with other pathway databases in enrichment analyses and machine learning. While PFOCR is applicable to any analysis involving pre-defined gene sets, we will highlight specific types of enrichment analyses and machine learning where its unique breadth and depth are crucial to fruitful results and insightful interpretation.

## Results

### Disease Coverage Comparison

A common goal of pathway analysis is to identify processes underlying a disease state. Thus, an important characteristic for a pathway database is its disease coverage, i.e., how many diseases are represented among its pathway-defined gene sets and how deep and diverse are the gene sets per disease. PFOCR has close to a hundred times as many pathways as a canonical pathway database, but how does their distribution across diseases compare to an intentionally curated database? In order to assess disease coverage, a standard set of 876 distinct disease names was compiled from the Comparative Toxicogenomics Database (CTD) [14]. The titles and descriptions associated with pathways from WikiPathways, Reactome, and KEGG were queried for disease name occurrences. Similarly, the titles and captions annotating PFOCR content were queried.

A total of 791 (90%) diseases were represented by at least one pathway in PFOCR. Reactome, WikiPathways and KEGG represented 153 (17%), 127 (14%), and 94 (11%) diseases, respectively (**Supplemental Table 1**). In terms of depth, we focused on the top 20 disease matches for each database; the union of these matches comprise the 37 diseases shown in **Table 1**. PFOCR includes 1954 pathways relevant to *breast cancer* and offers 649 pathways on average among its top 20 diseases. Reactome holds 23 pathways relevant to *leukemia* and 7 pathways on average among its top 20 diseases. WikiPathways has 18 pathways relevant to *SARS-CoV-2*—which aligns with its unique, community-driven approach to collecting new content on emerging topics—and also has 7 pathways on average among its top 20 diseases. KEGG has a four-way tie with 5 pathways for each of *lung cancer, insulin resistance, hepatitis*, and *cardiomyopathy*, and only 3 pathways on average among its top 20 diseases. By contrast, PFOCR has 5 or more pathways for 351 diseases and at least 1 pathway for 555 diseases that are missing from the other pathway databases (**Supplemental Table 1**).

**Table 1.** Disease Coverage Comparison. Union of top 20 disease-related pathways for each database by searching associated pathway titles and descriptions or captions. Disease names were compiled from the Comparative Toxicogenomics Database [14]. Average pathway counts among the top 20 disease matches per database are given in parentheses in column headers. The pathway count for the most represented disease per database is in bold italics. *Abbreviations: AML = acute myeloid leukemia, CDG = congenital disorders of glycosylation.

| Disease | PFOCR (649) | Reactome (7) | WikiPathways (7) | KEGG (3) |
| --- | --- | --- | --- | --- |
| Breast cancer | **1954** | 20 | 6 | 3 |
| Fibrosis | 959 | 4 | 8 | 4 |
| Alzheimer | 874 | 6 | 12 | 2 |
| Colorectal cancer | 817 | 10 | 5 | 1 |
| Prostate cancer | 794 | 1 | 7 | 1 |
| Lung cancer | 792 | 7 | 7 | **5** |
| Lymphoma | 692 | 9 | 2 | 3 |
| Leukemia | 673 | **23** | 6 | 3 |
| Melanoma | 659 | 8 | 4 | 1 |
| Insulin resistance | 586 | 1 | 4 | **5** |
| Obesity | 529 | 4 | 8 | 4 |
| Ischemia | 483 | 2 | 2 | 0 |
| Parkinson | 464 | 4 | 6 | 2 |
| Pancreatic cancer | 435 | 1 | 6 | 1 |
| Tuberous sclerosis | 413 | 1 | 0 | 0 |
| Hepatitis | 397 | 4 | 3 | **5** |
| Hypertrophy | 380 | 0 | 6 | 4 |
| Arthritis | 372 | 6 | 4 | 1 |
| Hypertension | 358 | 2 | 4 | 2 |
| Glioblastoma | 343 | 8 | 2 | 1 |
| Atherosclerosis | 300 | 6 | 4 | 3 |
| SARS-CoV-2 | 265 | 3 | **18** | 1 |
| Retinoblastoma | 204 | 3 | 5 | 2 |
| Heart failure | 185 | 0 | 2 | 3 |
| Anemia | 179 | 6 | 1 | 1 |
| Diabetes mellitus | 177 | 1 | 3 | 4 |
| Schizophrenia | 146 | 4 | 2 | 0 |
| Renal cell carcinoma | 142 | 0 | 5 | 1 |
| Cardiomyopathy | 133 | 1 | 1 | **5** |
| Huntington | 121 | 1 | 3 | 2 |
| Multiple myeloma | 120 | 4 | 0 | 0 |
| Hyperplasia | 100 | 2 | 2 | 3 |
| AML* | 96 | 6 | 1 | 1 |
| Ulcer | 67 | 1 | 0 | 3 |
| Hyperlipidemia | 33 | 0 | 6 | 0 |
| Hemophilia A | 3 | 4 | 0 | 0 |
| CDG* | 2 | 4 | 1 | 1 |

In order to assess gene coverage among disease pathways, we referenced the matching disease names in Jensen DISEASES, a database that provides gene-disease associations retrieved from text mining, literature, cancer mutation data, and genome-wide association studies [15]. **Table 2** shows these 17 matching diseases and the corresponding number of Jensen DISEASES genes covered by each pathway database. PFOCR covers 53 of cardiomyopathy genes (62%), which is the maximum count, and covers 63% of disease genes on average. Reactome has a maximum count of 12 (48%) for breast cancer and 9% on average. WikiPathways has a maximum count of 15 (19%) for diabetes mellitus and 21% on average. The maximum count by KEGG is for cardiomyopathy at 31 (36%) and the average coverage is 28%. For each of the diseases in **Table 2**, PFOCR covers a higher percentage of disease-associated genes than the other pathway databases. Interestingly, with the exception of diabetes mellitus, PFOCR includes over half of the genes associated with a given disease in pathway models labeled by those diseases.

**Table 2.** Disease Gene Coverage Comparison. Comparison of the disease-related gene content in the top disease-annotated pathways from Table 1 based on matching reference gene sets available from Jensen DISEASES. Absolute gene counts together with percentage of DISEASES genes in parentheses are provided for each database-disease pair. The gene counts for the most represented disease per database is in bold italics. An average percentage of DISEASES genes per database is given in parentheses in column headers.

| Disease | PFOCR (63%) | Reactome (9%) | WikiPathways (21%) | KEGG (28%) |
| --- | --- | --- | --- | --- |
| Cardiomyopathy | **53 (62%)** | 2 (2%) | 14 (16%) | **31 (36%)** |
| Diabetes mellitus | 31 (39%) | 2 (3%) | **15 (19%)** | 18 (23%) |
| Obesity | 45 (71%) | 4 (6%) | 8 (13%) | 12 (19%) |
| Parkinson | 34 (64%) | 3 (6%) | 11 (21%) | 15 (28%) |
| Prostate cancer | 23 (64%) | 2 (6%) | 6 (17%) | 5 (14%) |
| Alzheimer | 18 (58%) | 9 (29%) | 9 (29%) | 9 (29%) |
| Melanoma | 21 (70%) | 5 (17%) | 6 (20%) | 7 (23%) |
| Lung cancer | 24 (86%) | 7 (25%) | 13 (46%) | 17 (61%) |
| Breast cancer | 23 (92%) | **12 (48%)** | 14 (56%) | 13 (52%) |
| Hypertension | 7 (88%) | 2 (25%) | 3 (38%) | 3 (38%) |
| Atherosclerosis | 5 (63%) | 1 (13%) | 0 (0%) | 3 (38%) |
| Retinoblastoma | 2 (67%) | 0 (0%) | 1 (33%) | 1 (33%) |
| Huntington | 1 (50%) | 0 (0%) | 1 (50%) | 1 (50%) |
| Renal cell carcinoma | 1 (50%) | 0 (0%) | 0 (0%) | 0 (0%) |
| Schizophrenia | 8 (50%) | 1 (6%) | 1 (6%) | 0 (0%) |
| Multiple myeloma | 4 (50%) | 1 (13%) | 0 (0%) | 0 (0%) |
| Tuberous sclerosis | 2 (100%) | 0 (0%) | 0 (0%) | 0 (0%) |

### General Pathway Analysis

In practice, PFOCR is available in a format commonly used by enrichment analysis algorithms (see GMT in Data Availability). Thus, PFOCR can fit into analytical pipelines involving any package supporting GMT files, including the top-ranked Bioconductor packages: fgsea [16], clusterProfiler [17], GSEABase [18], and GSVA [19]. The pathway figure database has already been incorporated into online tools such as Enrichr[20] and NDEx iQuery [21].

Enrichr (https://maayanlab.cloud/Enrichr) provides quick and easy enrichment analysis against over 200 gene set databases simultaneously. Among the 27 databases in the pathway category, PFOCR has the greatest number of gene sets and the second largest coverage of human genes (the kinase co-expression gene sets by ARCHS4 has the largest). PFOCR ranks fourth among all 209 Enrichr databases in terms of gene set size. Among the suite of plots provided by the Enrichment Analysis Visualizer “appyter” for Enrichr databases [22], a UMAP helps to visualize the density and diversity of gene sets per database (**Figure 1A**). The appyter analysis identified 35 clusters of PFOCR pathways. By contrast, GO Biological Process divides into only 18 clusters, Reactome 27, WikiPathways 8, and KEGG 11, thus characterizing the greater diversity of pathway-based gene sets in PFOCR.

**Figure 1.**
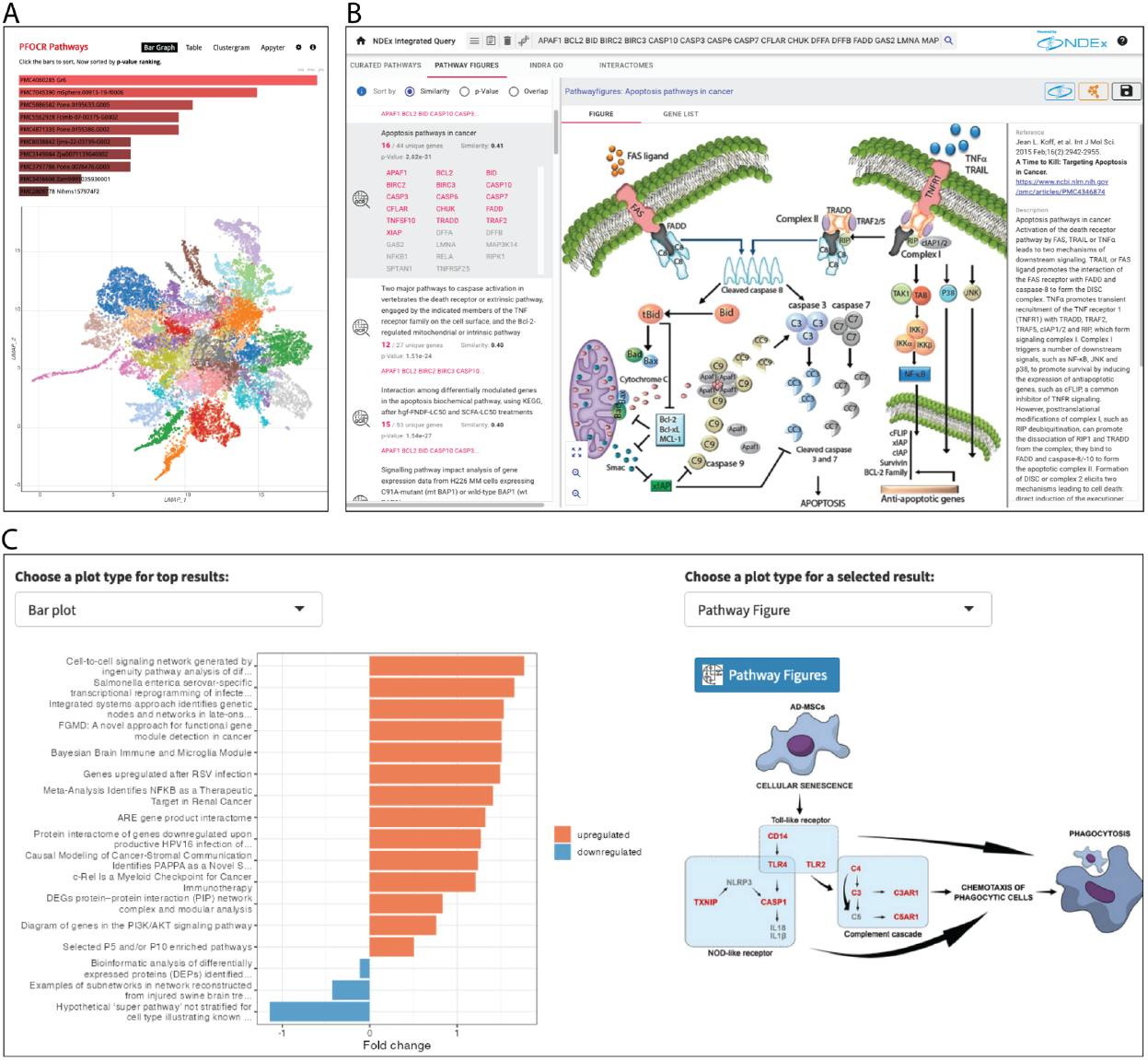
Pathway Analysis with PFOCR. (A) Typical bar graph of Enrichr results for PFOCR pathways, above. Appyter UMAP of PFOCR pathway clusters, below. The Enrichment Analysis Visualizer appyter computed term frequency-inverse document frequency (TF-IDF) values for the gene set corresponding to each pathway in PFOCR and plotted the first two dimensions of a UMAP applied to the resulting values. Generally, pathways with more similar gene sets are positioned closer together. Pathways are colored by Leiden algorithm-computed clusters. (B) NDEx iQuery screenshot showing ranked pathway figures (left panel), original pathway figure of selected result (middle), and article metadata (right). The top-right includes buttons to a dedicated NDEx page or to import the figure-extracted gene set into Cytoscape as nodes. (C) The plot options for ORA and GSEA results in the R Shiny tool called Interactive Enrichment Analysis. PFOCR results include a view of the original published figure and a link to a dedicated PFOCR website page (right).

NDEx Integrated Query (iQuery, https://www.ndexbio.org/iquery) provides network and pathway gene set analysis using multiple methods simultaneously against PFOCR, WikiPathways, INDRA-connected GO terms [23], and various interactomes [21]. Like Enrichr, the input is a simple list of gene names, and results are produced immediately. With a focus on networks and pathways, however, NDEx iQuery offers more detailed views of resulting gene sets ranked by similarity, p-value, or overlap (**Figure 1B**). In the case of PFOCR, the detailed view includes the original published figure next to its title, caption and other article metadata. Each pathway figure is associated with an interactive gene list and links to a dedicated page in the NDEx database. The PFOCR’s pathway figure-based gene sets can also be imported as nodes into Cytoscape with a single click.

We also developed an R Shiny tool called Interactive Enrichment Analysis (https://github.com/gladstone-institutes/Interactive-Enrichment-Analysis) to perform two types of enrichment analyses for one or more datasets simultaneously against GO, WikiPathways, and PFOCR [24]. The tool supports interactive exploration of results with customizable plots (volcano, dot, bar, heatmap, emap, and GSEA plots) and embedded pathway views (**Figure 1C**). In addition to views of original published figures, PFOCR results include links to dedicated web pages at the PFOCR database, which include a rich collection of metadata, crosslinks to PubMedCentral, NDEx and WikiPathways, and downloadable tables of extracted genes, chemicals and disease terms.

In the next two sections, we demonstrate advanced pathway analyses on two different disease datasets, tailoring methodology for each and demonstrating the level of interpretive detail that is only possible with a pathway database having PFOCR’s unique breadth and depth.

### Case Study 1: Acute Myeloid Leukemia Subtype Analysis

Leukemia is a fatal disease with a 5-year overall survival rate of 24% and a long-term survival rate of less than 20% in adulthood [25–28]. Among different types of leukemia, acute myeloid leukemia (AML) is characterized by clonal disorders of the hematopoietic compartment, such as abnormal proliferation of undifferentiated myeloid progenitors, impaired hematopoiesis, bone marrow failure and variable response to therapy [28]. Leukemia became a treatable disease with the development of drugs such as midostaurin, gilteritinib, and ivosidenib [29]. Studies have shown that leukemia drug efficacy is highly dependent on a patient’s genetic subtype profile. For example, midostaurin provides excellent treatment for patients with FLT3 mutations, whereas ivosidenib is particularly effective for patients with IDH1 mutations [30].

Additionally, Smoothened (SMO) inhibitors, such as Glasdegib, control the progression of acute leukemia by specifically targeting the Hedgehog (Hh) signaling pathway [31]. This subtype specificity can be effectively understood by investigating the effects of drugs on signaling cascades at the pathway level, rather than at a single gene level, assessing perturbations caused by different mutations [32]. Likewise, characterizing subtypes based on pathway-level transcriptomic profiles can help develop effective therapeutic strategies to optimize best survival outcomes of leukemia patients.

We characterized the perturbed transcriptomic profiles of AML subtypes using leukemia pathways from PFOCR, WikiPathways, Reactome, and KEGG. The purpose of the analysis was to compare the effectiveness of these pathway databases in characterizing leukemia subtypes based on gene expression. AML patient gene expression data with 8 mutations was retrieved from GEO (GSE108316) and PCA-based quality control was performed. We then performed differentially expressed gene (DEG) analysis, where each mutation type was treated as a separate group and control samples were used as the reference, followed by gene set enrichment analysis (GSEA). Finally, hierarchical clustering was applied to the normalized enrichment scores (NES) from GSEA to cluster leukemia mutations in operational subtypes.

A total of 705 leukemia-enriched pathways were investigated by hierarchical clustering analysis, including 673 PFOCR, 6 WikiPathways, 23 Reactome, and 3 KEGG pathways (**Figure 2A**). At the top level of the hierarchical cluster, there were two core subclusters where the larger cluster (subtype L) was defined by five mutations, RUNX1, INV16, t(8:21), CEBPA, and SRSF2, and the smaller cluster (subtype S) was defined by three mutations, FLT3-ITD, FLT3-ITD/NPM1, and inv3/RAS. Among all mutations of subtype S, FLT3 mutation is associated with the most unfavorable prognosis [29,34]. FLT3, a transmembrane ligand-activated receptor tyrosine kinase, is expressed in hematopoietic progenitor cells, and 25-30% of AML cases carry FLT3 mutations that result in abnormal cell growth and survival via mTOR and PI3K/AKT pathway [35]. In subtype L, RUNX1 is another well-known type of AML mutation associated with hematopoietic stem cell (HSC) growth, differentiation, and homeostasis. Its abnormalities are often associated with older age and male sex, and are found in 8–16% of AML patients [35].

**Figure 2.**
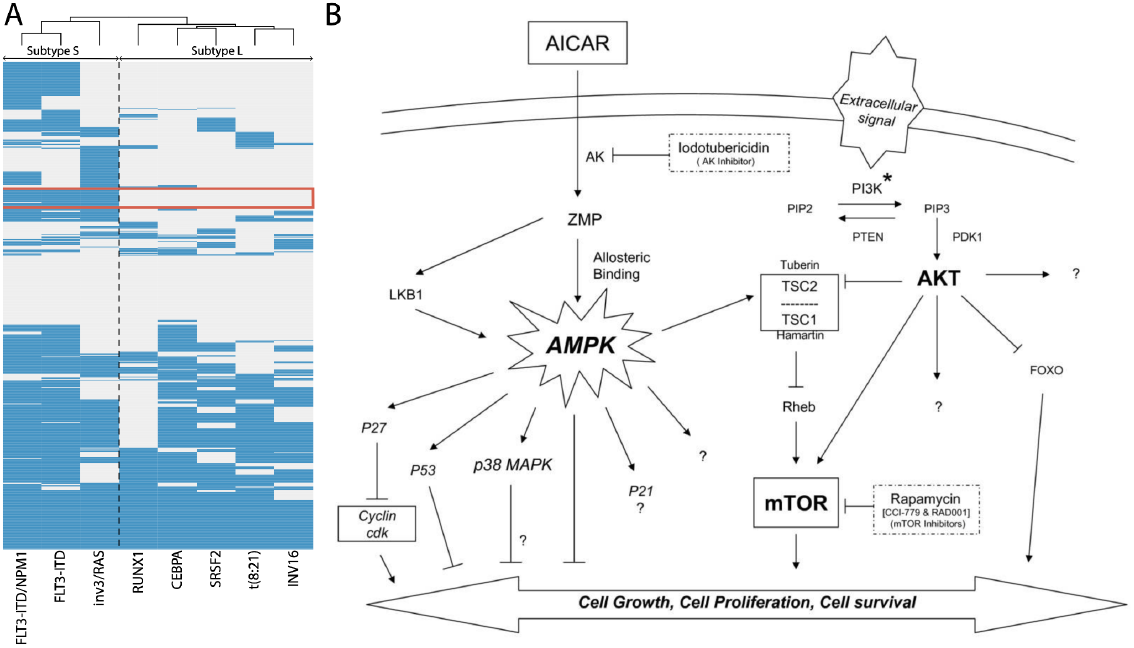
Heatmap of GSEA NES values from leukemia expression data. (A) Hierarchical clustering of GSEA NES values of leukemia mutations defines two top-level subtypes, denoted S and L. The cluster of 26 pathways that differentiates the two subtypes is highlighted with a red box. (B) PFOCR’s PMC1948012 F10 [33] is one of the 26 subtype-distinguishing pathways and contains the FOXO gene signaling pathway. This figure is reproduced from [28].

Of the 705 leukemia pathways, there are 26 pathways that clearly distinguish subtypes L and S (**Figure 2A, boxed rows, Supplemental Table 2**). These 26 core leukemia gene sets (CLGS) consist of 24 PFOCR pathways, 1 WikiPathways and 1 KEGG pathway. The predominance of PFOCR pathways in defining the CLGS highlights the utility of PFOCR in characterizing subtypes compared to other pathway databases. For example, one of the PFOCR pathways in the CLGS involves the regulation of forkhead box (FOX) genes through PI3K/AKT signaling cascade (**Figure 2B**). FOX genes are transcription factors involved in the regulation of multiple cellular functions, including development, differentiation, proliferation, and apoptosis [36–38], and may act as either tumor suppressors or oncogenes depending on the cellular and biological context [28,39]. In the context of leukemia, FOX genes have been reported to play different roles depending on the type of mutation. For example, upregulation of FOXO1 in AML with a RUNX1 mutation has been reported to help the growth of leukemia cells and inhibit the differentiation of CD34+ hematopoietic stem and progenitor cells [28]. In addition, the activity of FOXO1 has been reported to affect the antineoplastic drug sensitivity of AML cells, and has been proposed as a therapeutic for leukemia with RUNX1 mutations [40,41]. On the other hand, FOXO3 acts as a tumor suppressor, and phosphorylation of FOXO3 by FLT3-ITD results in inactivation of FOXO3-mediated apoptosis in leukemia with FLT3-ITD mutation [42].

As mentioned above, the success of FDA-approved leukemia drugs depends on a patient’s tumor genetics. For example, a study tested the effect of midostaurin in leukemia patients with RUNX1 showed a limited role of the drug in leukemia control [43]. Midostaurin treatment in systemic mastocytosis patients has shown that patients with one or more mutations in the S/A/R (SRSF2, ASXL, or RUNX1) panel have a lower survival rate and a higher progression rate to AML or mast cell leukemia than patients without mutations. Additionally, the same study found that midostaurin treatment was not able to prevent the increase of RUNX1 mutations in patients which was associated with progression to secondary AML [43]. To date, there have been no officially approved drugs effective for leukemia with RUNX1 mutations [29,34,44].

The different roles of FOXO genes in RUNX1 leukemia and FLT3-ITD leukemia highlights the mechanistic differences between the two leukemia subtypes and why treatment of both subtypes with the same drug may fail. PFOCR-based gene set hierarchical clustering analysis is an effective methodology that can aid in understanding pathway-level mechanisms underlying the differences between cancer subtypes.

### Case Study 2: Breast Cancer Prediction Analysis

To evaluate the disease prediction efficacy of these pathway databases, machine learning analysis was performed on breast cancer patient expression data. For this analysis, two independent breast cancer patient expression data sets were collected from GSE3494 and GSE2990, and samples with breast cancer grades 1 and 2 were selected for analysis. 51 samples with a grade value of 2 were randomly selected from GSE3494 to balance the number of samples in both grades. For the test data, GSE2990 was used, and there were 29 grade 1 and 31 grade 2 samples. Random forest was chosen to build predictive models, and tunned model parameters were selected based on out-of-sample error. On all pathways annotated as breast cancer, prediction accuracy was calculated from the test data and leave-one-out cross-validation accuracy was calculated from the training data. Gene importance (feature importance) was calculated as the average importance from the leave-one-out cross-validation iterations. The prediction accuracy rankings in **Table 3** shows 21 pathways with prediction accuracy and cross-validation accuracy larger than 0.65 (bold), which are all from PFOCR. **Supplemental Table 3** presents all results with prediction accuracy and cross-validation accuracy larger than 0.55 and min gene importance larger than 0.1, which include 695 PFOCR, 9 Reactome, and 1 KEGG pathways.

**Table 3.** Random forest breast cancer grade prediction accuracy with top important gene information. Pathways with cross-validation accuracy and prediction accuracy greater than 0.65 (all from PFOCR), plus the top ranked results from Reactome and KEGG (discontinuously below dashed line). The result ranking was determined by max(min(cross-validation accuracy, prediction accuracy)). The “Top Gene” corresponds to the gene with the highest feature importance score in each pathway. The “Ref.” column provides paper and pathway citations for each result.

| Rank | Pathway | Ref. | Top Gene | Importance Score | CV Accuracy | Prediction Accuracy | Specificity | Sensitivity | F1 |
| --- | --- | --- | --- | --- | --- | --- | --- | --- | --- |
| 1 | PMC2937358_F1 | [45,46] | CXCL6 | 0.61 | 0.72 | 0.72 | 0.61 | 0.83 | 0.69 |
| 2 | PMC2653381_F1 | [47,48] | GSTM4 | 0.25 | 0.69 | 0.7 | 0.58 | 0.83 | 0.67 |
| 3 | PMC5772637_F6 | [49,50] | CASP9 | 0.91 | 0.69 | 0.68 | 0.84 | 0.52 | 0.73 |
| 4 | PMC5772637_F7 | [49,51] | CASP9 | 1.54 | 0.68 | 0.72 | 0.77 | 0.66 | 0.74 |
| 5 | PMC8040471_F7 | [52,53] | CASP9 | 0.45 | 0.68 | 0.67 | 0.81 | 0.52 | 0.71 |
| 6 | PMC6759650_F5 | [54,55] | CYCS | 0.37 | 0.67 | 0.78 | 0.71 | 0.86 | 0.77 |
| 7 | PMC7811378_F9 | [56,57] | CASP9 | 0.85 | 0.67 | 0.78 | 0.84 | 0.72 | 0.8 |
| 8 | PMC5715135_F7 | [58,59] | CASP9 | 0.15 | 0.67 | 0.75 | 0.84 | 0.66 | 0.78 |
| 9 | PMC2673236_F1 | [60,61] | ACSM3 | 1.32 | 0.67 | 0.72 | 0.74 | 0.69 | 0.73 |
| 10 | PMC387764_F8 | [62,63] | CASP9 | 1.78 | 0.67 | 0.68 | 0.74 | 0.62 | 0.71 |
| 11 | PMC6947643_F2 | [64,65] | HEY1 | 1.23 | 0.67 | 0.67 | 0.84 | 0.48 | 0.72 |
| 12 | PMC7409684_F1 | [66,67] | WNT11 | 0.18 | 0.67 | 0.67 | 0.68 | 0.66 | 0.68 |
| 13 | PMC8023395_F2 | [68,69] | EGFR | 0.17 | 0.66 | 0.73 | 0.84 | 0.62 | 0.76 |
| 14 | PMC4336604_F9 | [70,71] | MAPK9 | 0.33 | 0.66 | 0.72 | 0.74 | 0.69 | 0.73 |
| 15 | PMC6499473_F1 | [72,73] | PARP1 | 0.23 | 0.66 | 0.72 | 0.61 | 0.83 | 0.69 |
| 16 | PMC3219187_F5 | [74,75] | ZBTB7A | 2.29 | 0.66 | 0.7 | 0.81 | 0.59 | 0.74 |
| 17 | PMC6024909_F4 | [76,77] | CDK1 | 0.26 | 0.66 | 0.7 | 0.52 | 0.9 | 0.64 |
| 18 | PMC2694962_F1 | [78,79] | ACSM3 | 1.52 | 0.66 | 0.68 | 0.48 | 0.9 | 0.61 |
| 19 | PMC5256616_F6 | [80,81] | CASP9 | 0.36 | 0.66 | 0.68 | 0.48 | 0.9 | 0.61 |
| 20 | PMC4407294_F2 | [82,83] | NOTCH2 | 0.7 | 0.66 | 0.67 | 0.52 | 0.83 | 0.62 |
| 21 | PMC6305585_F2 | [84,85] | CPEB1 | 0.77 | 0.66 | 0.67 | 0.71 | 0.62 | 0.69 |
| 63 | R-HSA-8864260 | [86] | HSPD1 | 0.25 | 0.64 | 0.68 | 0.81 | 0.55 | 0.72 |
| 115 | KEGG_hsa01522 | [87] | BAX | 0.11 | 0.62 | 0.85 | 0.9 | 0.79 | 0.86 |

To assess the overlap of gene content among the top 21 pathways from PFOCR and any of the results from the other databases that passed the minimum accuracy 0.55 threshold, we plotted the genes per pathway, ordered (and sized) by their importance scores (**Figure 3**). Remarkably, the majority of genes, including top scoring “important” genes, from the highest accuracy pathways are unique to PFOCR results.

**Figure 3.**
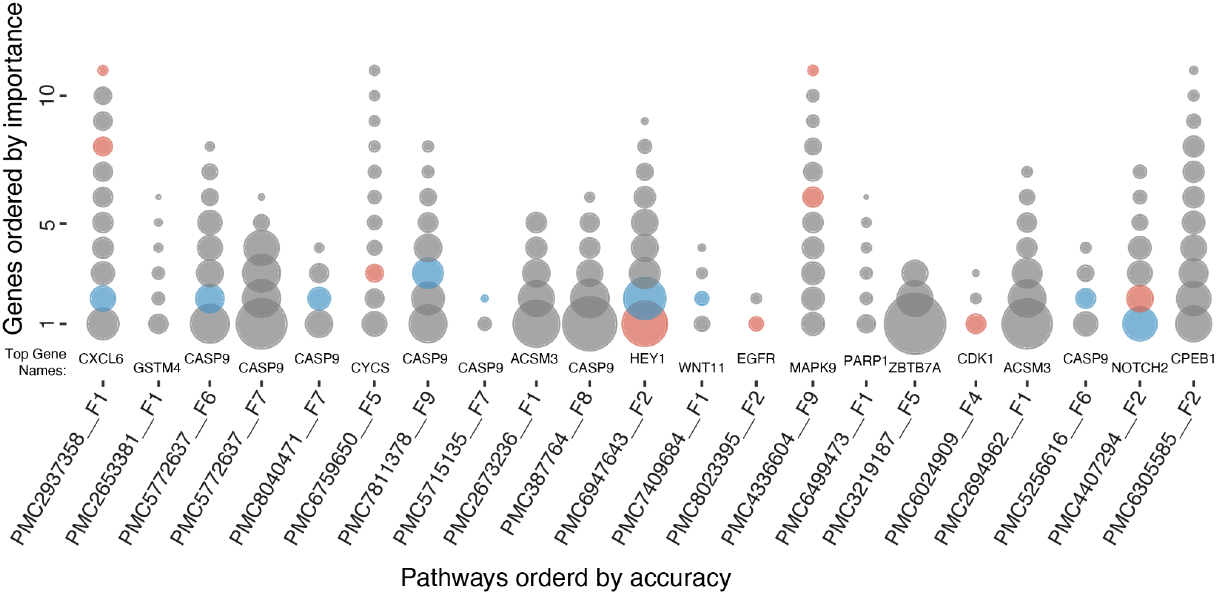
Gene importance from random forest breast cancer models built on pathway genes. Represented as circles, the gene content is sized and ordered by importance (large-to-small, bottom-to-top) for the top twenty-one pathways ordered by prediction accuracy (left-to-right). The “Top Gene” from Table 3 is represented by the largest, bottom-most circle and is labeled for each pathway. The vast majority of circles (gray) denote genes found only in PFOCR pathways among all significant pathways with “important” genes. The genes that are also present within Reactome and KEGG results are highlighted in red and blue, respectively. There were no significant pathways with gene importance scores above 0.1 from WikiPathways.

CASP9, the gene with the second highest importance score in 21 PFOCR pathways, is an initiator of apoptosis in the mitochondrial apoptosis pathway. A study by Sharifi and Moridnia found an association of CASP9 expression with miR-182-5p [88]. The breast cancer cell line MCF-7 had poor viability when miR-182-5p was inhibited, suggesting CASP9 upregulation is related to MCF-7 cell viability. In addition, an independent study investigated SNPs in CASP9 to find an increased breast cancer risk in patients with CASP9 mutations. In particular, CASP9 SNPs rs4645978 and rs4645981 were associated with high breast cancer risk, suggesting that CASP9 contributes to breast cancer development [89].

Among the overlapping genes of PFOCR and REACTOME, HEY1, one of the well-known Notch target genes, had the highest shared gene importance score. Chen et al. investigated the expression of HEY1 in breast cancer cells and found that the HEY1 expression level increased under hypoxic conditions [90]. In addition, increased Notch4-Hey1 mRNA expression and decreased patient survival were found to be correlated in another study, confirming Hey1 as a marker for breast cancer development [91].

On the other hand, NOTCH2 was the gene with the highest importance score overlapping between PFOCR and KEGG. The role of NOTCH2 was studied by Fu et al. by investigating NOTCH2 expression and polymorphisms of SNP rs11249433 in breast cancer patient data [92]. That study suggested that increased expression of NOTCH2 with the rs11249433 polymorphism may contribute to the development of ER+ luminal tumors.

## Discussion

Using the Comparative Toxicogenomics Database (CTD) [14] for a comprehensive, independent set of 876 disease names, we chose the most comparable sources of metadata (titles and descriptions or captions) to assess the disease coverage by PFOCR and popular pathway databases: Reactome, WikiPathways, and KEGG. In terms of both breadth and depth, PFOCR surpassed the other pathway databases by a considerable margin. We assessed the most common diseases represented by each database (e.g., PFOCR has 1954 breast cancer pathways), the average number of pathways among their top 20 diseases (e.g., PFOCR has 649 pathways on average), and the minimal coverage (e.g., PFOCR has 5 or more pathways for 351 diseases and at least 1 pathway for 555 diseases that are lacking representation from any of the other pathway databases). In terms of actual disease gene content in these pathways, we made use of Jensen DISEASE [15] annotations and focused on 17 diseases including the best representatives from each database and spanning neurological, cancer, heart, lung and metabolic categories. Again, the contrast was striking, with PFOCR covering more disease genes in every case and over half of the genes in all diseases except diabetes mellitus (39%), while the other pathway databases averaged between 9% and 28% coverage. Researchers interested in associating their genes of interest (e.g., from differential gene expression analyses) with mechanisms of disease will find broader coverage and more diverse instances in the PFOCR database.

Like other pathway databases, PFOCR content is available in formats amenable to gene set enrichment analyses, such as GMT (see Data Availability), allowing bioinformaticians to include PFOCR in practically any pathway analysis workflow. PFOCR pathways have already been integrated into commonly used online tools for enrichment analyses, including Enrichr [20] and iQuery by NDEx [21], and we also have introduced an R Shiny app called Interactive Enrichment Analysis, which uses PFOCR, WikiPathways and GO by default [24].

AML subtype clustering analysis showed how PFOCR pathway information can be used to gain insights to understand differential regulatory mechanisms of disease subtypes at the pathway level. One example of a more general approach of enrichment analysis using canonical gene sets was demonstrated in the study performed by Asi et al. [93]. In this study, enrichment analysis was performed using KEGG and GO on the combined data of DNase I-seq and RNA-seq data from FLT3-ITD and t(8;21) AML patient samples. Integration was performed by finding DNase I hypersensitivity site (DHS) peaks specific to two mutation types and combining the result with the DEG analysis result. Subsequently, GO and KEGG enrichment analysis performed for each subtype and enriched gene sets for each AML subtype were investigated. This study provided a list of gene sets and compared the presence/absence of gene set signals in each subtype. While this approach is the most common way gene set enrichment is used in research, utilizing PFOCR with its extensive collection of more diverse gene sets and contexts can help to investigate pathway level signals in greater detail. We also demonstrated how clustering approaches can make gene set comparisons more systematic and interpretable by categorizing pathway-level signals specific to each subtype.

Each published pathway figure offers not only unique content, but also unique context based on the specific experiments and insights described in its parent article. As a collection, PFOCR offers a distinct advantage to researchers in providing diverse examples of a given biological process linked to specific experimental designs and analytical methods. For example, our case study on breast cancer grade prediction showed that PFOCR outperformed other pathway databases in terms of both rank and total number of results. In our breast cancer analysis results, the top 62 pathways that passed the accuracy threshold were all from PFOCR. The fact that the majority of genes—including those with the highest importance scores—were unique to PFOCR suggests that the unique pathway information in PFOCR can support research into disease mechanisms in a way that other pathway databases cannot.

## Conclusions

PFOCR is a novel kind of pathway database approaching the breadth and depth of GO while providing rich, mechanistic diagrams and direct literature support. PFOCR content is extracted from the pathway figures embedded in published articles dating as far back as 1995 and at a current pace of one thousand new pathways each month. Compared to other popular pathway databases, PFOCR contains an unprecedented number and diversity of pathways, genes, and chemicals. Another 23,000 pathways figures will be added to the database soon, covering 2021-2023, with plans to regularly update as new figures are published. With a focus on disease-associated content, PFOCR leads not only in terms of the number of diseases covered, but also the number of pathways per disease, and the number of unique genes per disease. In the context of pathway analysis, PFOCR offers more avenues to connect a researcher’s gene set of interest to biological processes related to disease. Available in GMT format and pre-integrated into user-friendly online tools, PFOCR is easy to include in a researcher’s analysis plan. More advanced pathway analyses, for example investigating cancer subtypes and grade prediction, can leverage the unique depth of pathway content in PFOCR that lends support to explorations into possible mechanistic models and machine learning applications where typical pathway databases are not particularly useful.

## Methods

The PFOCR gmt (pfocr-20210515-gmt-Homo_sapiens.gmt) includes figures published between 1995 and 2021 and was downloaded from the PFOCR data archive [94]. The WikiPathways gmt file (wikipathways-20211210-gmt-Homo_sapiens.gmt) includes curated pathway models up until 2021 and was downloaded from the WikiPathways data archive [95]. KEGG pathways were retrieved from json files downloaded from TogoWS [96]. TogoWS only supports KEGG data uploaded prior to their proprietary licensing in 2012. Reactome pathways were retrieved as JSON files using the Reactome REST API on 10 January 2022, and were parsed using the R jsonlite library [97].

PFOCR, Reactome, WikiPathways, and KEGG pathways are filtered according to the criteria of having a minimum of 3 genes and a maximum of 500 genes. In order to best represent Reactome’s unique gene-level annotations, no gene minimum was applied for the disease gene coverage comparison (**Table 2**). As a result of these filters, the number of pathways from their sources decreased from 2029 to 1663 in Reactome, from 703 to 686 in WikiPathways, and from 345 to 345 in KEGG. The PFOCR gmt is provided with these constraints already applied.

### Disease coverage comparison

Disease information was downloaded from the Comparative Toxicogenomics Database (CTD) [14]. Diseases were screened according to the following criteria in order to compile a distinct set of names amenable to making unambiguous occurrence counts in text annotations in pathway databases: 1. A disease name is not an extension of another disease name from the same disease, 2. A disease name is not related to a psychological condition, 3. A disease is not a category for multiple diseases otherwise included (e.g., neurodegenerative disease), 4. A disease is not related to an environmental condition (e.g., mite infestations), 5. A disease is not an alias for another disease, 6. A disease is not a symptom (e.g., abdominal pain), 7. A disease name is not ambiguous relative to included disease names (e.g., cancer). The final number of filtered disease names was 876. Text titles and descriptions (or captions) were collected for each of the human pathways from PFOCR, Reactome, WikiPathways, and KEGG. Case-insensitive string matching functions were used to identify disease name occurrences in the collected text samples. A match was only counted once per pathway even if the disease name occurred multiple times within or across text samples for that pathway. The resulting pathway counts per disease and per database are shown in **Supplement Table 1** and a subset in Table 1. Reactome and WikiPathways provide additional sources for disease annotation, including ontology tags, gene descriptions, and bibliography titles that we did not include in this accounting in order to make a fair comparison across all four resources.

To investigate disease gene coverage of the pathway databases, the human disease gene file ‘human_disease_knowledge_filtered.tsv’ was downloaded from Jensen DISEASES [98]. Jensen disease names that exactly matched the CTD disease names were selected for investigation. The number of genes present in integrated pathways for each disease was determined for each pathway database and also expressed as a percentage of the number of genes defined by Jensen DISEASES for each disease.

### Case Study 1: Acute Myeloid Leukemia Subtype Analysis

AML patient gene expression data was downloaded from GEO under accession number GSE108316.

Samples with mutations in RUNX1, inv(16), t(8;21), CEBPA, SRSF2, FLT3-ITD, FLT3-ITD/NMP1, and inv(3)/RAS were selected for this study. Differentially expressed gene (DEG) analysis was performed using genes with two or more read counts in at least five samples using the limma pipeline [99]. Gene counts were transformed to log_2_-counts per million (logCPM), and the mean-variance relationship and weights were estimated using voom in the R library limma [99]. Then, limma’s lmFit was used to fit linear models and limma’s eBayes to compute statistics of the fitted models. Control samples were used as reference samples for each hypothesis comparing leukemia patients and normal gene expression levels.

Based on the DEG results, GSEA was performed for each mutation type. First, leukemia related pathways were retrieved from PFOCR, WikiPathways, KEGG, and Reactome (see eighth row of **Table 1**). GSEA was performed using GSEA in the R library ClusterProfiler [17]. Normalized enrichment scores (NES) were retrieved from GSEA results, and scores for each mutation type were subjected to hierarchical clustering using the R function *heatmap*.*2*. At the top level of the sample hierarchy, there are two main clusters. The core leukemia gene set (CLGS) was defined by the hierarchical clustering and included 26 pathways with the greatest average NES differences.

### Case Study 2: Breast Cancer Prediction Analysis

GSE3494 and GSE2990 breast cancer patient gene expression data measured by Affymetrix Human Genome U133A Array were downloaded for breast cancer analysis. Robust Multi-array Average (RMA) [100] was calculated using rma in the R library affy. Principal component analysis (PCA) was performed to confirm that there was no batch effect in data sets.

To build random forest models for predicting breast cancer grade in patient data, first, breast cancer pathways were retrieved from PFOCR, Reactome, WikiPathways, and KEGG. Breast cancer pathways compiled from the disease coverage comparison analysis resulted in 1954 PFOCR, 20 Reactomes, 6 WikiPathways and 3 KEGG pathways (see first row of **Table 1**). The gene expression values of genes in each pathway were used as feature values in random forest models. For best results, hyperparameters (number of features, minimum node size, and fraction of observations to sample) were tuned using the tune function in the R library randomForestSRC [101] to ensure that the prediction error is the minimum. The number of trees was set to 1000 times the number of features. Then, ranger [102] was used to make random forest models and measure feature importance given the optimal hyperparameters. The overall training performance of each model and the feature importance was measured as the average of the one-out cross-validation results. The pathways with cross validation accuracy and prediction accuracy higher than 0.65 were selected as top pathways (**Table 3**) and pathways with cross validation accuracy and prediction accuracy higher than 0.55 were selected (**Supplemental Table 3**) and used to assess the uniqueness of top genes in PFOCR pathways.

## Declarations

### Availability of data and materials

All materials are available online. See website links and references.

### Authors’ contributions

All the analyses were performed by M-GS. Figures and tables were prepared by M-GS and ARP. The manuscript was written by M-GS and ARP.

## Acknowledgements

We would like to acknowledge Reuben Thomas, Martina Summer-Kutmon, Kristina Hanspers, Anders Riutta, and Laurent Winckers for early discussions and brainstorming leading to the conception of this manuscript.

